# Substrate availability and citrate alter TCA cycle metabolism and SLC13A3 in macrophage immune responses

**DOI:** 10.1101/2025.09.22.677838

**Authors:** Lea Woyciechowski, Hanna F. Willenbockel, Thekla Cordes

## Abstract

Cell culture media are commonly formulated to enhance cell growth and often lack the physiological nutrient composition found in human blood plasma. The impact of substrate availability on immune cell metabolism and function remains incompletely understood. Here, we demonstrate that changes in culture medium composition affect mitochondrial metabolic pathways, immune responses, and transport in macrophages. Using mass spectrometry and stable isotope tracing, we identify citrate as a mediator linking extracellular substrate availability to intracellular metabolism. We also observe increased IL-6 secretion and elevated expression of plasma membrane transporter NaDC3 (SLC13A3) under physiological carbon source conditions that are reversed when citrate is excluded from the medium. Our findings demonstrate that extracellular substrate composition shapes macrophage immunometabolism and identify citrate as an extracellular signal that modulates immune responses. This work highlights the importance of physiologically relevant nutrient availability in studying and targeting immunometabolic pathways.

## Introduction

In the 1950s, the development of cell culture media was primarily focused on supporting cell proliferation and survival in vitro^1,2^. Among them, Dulbecco’s Modified Eagle’s Medium (DMEM) is widely used for mammalian cell culture. However, the media composition differs from that of physiological plasma and varies in nutrient and carbon source concentrations^3^. Several metabolites present in plasma, such as alanine, asparagine, citrate, lactate, pyruvate, and β-hydroxybutyrate (βHB), are absent from standard DMEM despite their biological function. For instance, lactate was once considered a metabolic end product of glycolysis and is now appreciated as a carbon fuel and regulator of redox balance^4–7^. While nutrient availability affects cellular metabolism and function^8–10^, the mechanisms by which altered substrates influence immunometabolism are incompletely understood.

Macrophages sense cues from their microenvironment and metabolic reprogramming is a hallmark of the innate immune response^11^. Mitochondria play a central role in orchestrating these responses and tricarboxylic acid (TCA) cycle intermediates act as signaling molecules to alter biological processes^12–14^. Itaconate is catalyzed via immunoresponsive gene 1 protein (IRG1/ACOD1) which is encoded by immune responsive gene 1 (*Irg1*) and has emerged as a key immunomodulatory effector molecule^15–17^. This molecule influences diverse metabolic pathways including acting as a competitive inhibitor for succinate dehydrogenase (SDH)^18–20^. Since itaconate is derived from TCA cycle intermediate cis-aconitate, changes in the availability of carbon sources for mitochondrial metabolism may affect itaconate synthesis.

Stable isotope tracing and mass spectrometry techniques help to reveal how immune cells reprogram their metabolism and use alternative substrates^21,22^. In addition to glucose and glutamine, alternative carbon sources such as lactate, specific amino acids, and ketone bodies provide fuel for the TCA cycle and support the synthesis of citrate^7,23–25^. The solute carrier (SLC) family is the largest group of transporters that facilitate the import of key metabolites across the plasma membrane. Among them, the sodium-dependent dicarboxylate transporter 3 (NaDC3/SLC13A3) transports TCA cycle-related metabolites, such as citrate, α-ketoglutarate (αKG), succinate, and malate. More recently, this carrier has been identified as an itaconate transporter^26–28^. However, how physiological carbon availabilities modulate immune cell metabolism, fueling TCA cycle metabolism, and how NaDC3 responds to changes in the extracellular environment remains incompletely understood.

Here, we developed a modified cell culture medium (mDMEM) to more closely recapitulate physiological carbon sources present in plasma. We applied mass spectrometry and functional assays to interrogate how the presence of alternative carbon substrates influences mitochondrial TCA cycle metabolism, itaconate synthesis, and immune responses in murine RAW264.7 macrophage-like cells. We demonstrate that citrate acts as an immunomodulatory metabolite in mDMEM cultures altering *Slc13a3* expression. Our findings highlight how media composition influences immunometabolism with potential implications for therapeutic strategies.

## Material and Methods

### Cell lines

The murine macrophage-like cell line RAW264.7 (TIB-71, ATCC) were cultured in high glucose Dulbecco’s Modified Eagle Medium (DMEM, Cat.# 41965-039, Gibco), supplemented with 50 U/ml penicillin, 50 µg/ml streptomycin (P/S, Cat.# 15140-122, Gibco) and 10 % (v/v) Fetal Bovine Serum (FBS, FBS.SAM.0500, Bio&Cel, sourced from South America) at 37 °C with 5 % CO_2_ and 21 % oxygen in a humidified incubator. Huh7 cells were detached with 0.05 % trypsin-EDTA (Cat.# 25300-054, Gibco). Cells were tested negative for mycoplasma contamination using MycoAlert® Mycoplasma Detection Kit (Cat.# LT07-118, Lonza).

### Cell treatment

The experiments analyzing metabolism and gene expression of the cells were performed in 12-well cell culture plates. RAW264.7 macrophages were seeded with a density of 3 × 10^5^ cells/well. The workflow involved a resting phase for 3 to 4 h post-seeding followed by an adaptation period of 36 h, during which a medium change to the adapted medium was performed around 24 h after the first medium change. Approximately 48 h after post-seeding, the cells were treated and stimulated with 10 ng/ml lipopolysaccharide (LPS, stock 1 mg/ml, *Escherichia coli* O128:B12, L2755, Sigma-Aldrich) for the RAW264.7 macrophages. Harvesting of the cells or the corresponding assay was performed after a 6 h incubation period with LPS.

### Isotopic tracing and analysis

Tracing of RAW264.7 cells was performed in DMEM (Cat.# D5030, Sigma-Aldrich) supplemented with 3.7 g/l bicarbonate. For ^13^C glucose tracing experiments, unlabeled glucose was replaced with 25 mM [U-^13^C_6_]glucose (Cat.# CLM-1396-1, Cambridge Isotope Laboratories). For ^13^C glutamine tracing experiments, unlabeled glutamine was replaced with 4 mM [U-^13^C_5_]glutamine (Cat.# CLM-1822-H-0.5, Cambridge Isotope Laboratories). All tracer media were sterile-filtered through 0.22 µm filters and supplemented with 10 % FBS, 50 U/ml penicillin, and 50 µg/ml streptomycin. For inflammatory stimulation, cells were stimulated with LPS as indicated in each figure legend. The [2,4-^13^C_2_]citrate tracer (Cat.# 492078, Sigma Aldrich) was added at a concentration of 0.215 mM to the growth media. Labeling on metabolites was quantified using gas chromatography coupled to mass spectrometry (GC-MS).

### Metabolite extraction

Metabolites were extracted as previously described in detail^29^. Briefly, cells were washed with saline solution (0.9 % w/v NaCl) and quenched with 0.25 ml -20 °C methanol. After adding 0.1 ml 4°C cold water containing an internal standard norvaline (5 µg/ml). The mixture was collected into a pre-cooled microcentrifuge tube and 20 µl of the methanol-water mixture was transferred into a 96-well plate for protein quantification. Next, 0.25 ml -20 °C chloroform was added to the sample, and the extracts were vortexed for 10 min at 4 °C and centrifuged at 16,000 × *g* for 5 min at 4°C. The upper aqueous phase was evaporated under vacuum at 4 °C and used for GC-MS analysis. For the medium samples, 10 µl of the medium was used for metabolite extraction and mixed with 80 µl of extraction fluid (methanol:H_2_O is 9:1), vortexed for 15 sec, and centrifuged at 17,000 x *g* for 5 min at 4°C. 80 µl of the upper phase were transferred to an analytical vial for the metabolite derivatization and evaporated.

### Gas Chromatograph–Mass Spectrometry (GC-MS) and sample preparation

Metabolites were analyzed and quantified, as previously described^29^. Polar metabolites were derivatized using a Gerstel MPS system with 15 μl of 2% (w/v) methoxyamine hydrochloride (Sigma-Aldrich) in pyridine (incubated for 90 min at 55°C) and 15 μl N-tertbutyldimethylsilyl-N-methyltrifluoroacetamide (MTBSTFA) with 1% tert-butyldimethylchlorosilane (tBDMS) (RESTEK, Cat.# 35601) incubated for 60 min at 55°C. Derivatives were analyzed by gas chromatography (GC) mass spectrometry (MS) using a ZB-35MS column (Phenomenex, Cat.# 7HG-G003-11-GGA-C) installed in an Agilent 7890B gas chromatograph interfaced with an Agilent 5977B MSD mass spectrometer operating under electron impact ionization at 70 eV. The MS source was held at 230 °C, the quadrupole at 150 °C, and helium was used as a carrier gas. The GC oven temperature was held at 100 °C for 2 min, increased to 325 °C at a rate of 10 °C/min, and held at 325 °C for 4 min. The metabolite identification of the GC-MS data was performed by using open chrome software (Lablicate) to determine the retention times for each metabolite. Mass isotopomer distributions and total metabolite abundances were computed by integrating mass fragments with corrections for natural isotope abundances using in-house algorithms^29^. Metabolite abundances were normalized to total cell protein content.

### Respirometry

The Seahorse assay was performed following the manufacturer’s instructions for the Agilent Seahorse XF Cell Mito Stress Test Kit (Agilent Technologies). Cells were seeded into a 96-well Seahorse plate at a density of 2 × 10^4^ cells/well with six cellular replicates per condition. For the assay medium, DMEM 5030 without glucose, L-glutamine, phenol red, sodium pyruvate, and sodium bicarbonate was used. For the regular medium 2 mM HEPES, 8 mM glucose, 2 mM glutamine, and 2 mM sodium pyruvate was supplemented and for the adapted medium 2 mM HEPES, 5 mM glucose, 0.5 mM glutamine, 0.15 mM sodium pyruvate, 0.19 mM asparagine, 0.4 mM alanine, and 0.4 mM acetate, 0.4 mM alanine, 0.19 mM asparagine, 0.215 mM citrate, 3 mM lactate, and 0.85 mM β-hydroxybutyrate (βHB).

Following the completion of the treatment period, the medium of the cell culture plate was changed to the assay medium. The cells were washed twice with 100 µl of the assay medium. Afterward, 150 µl of the assay medium was added to each well. The cell culture plate was incubated for approximately 20 min at 37 °C without CO_2_. Meanwhile, 25 µl of the inhibitors were added to designated ports. Port A contained oligomycin A [14 µM], while port B and C contained Trifluoromethoxy carbonylcyanide phenylhydrazone (FCCP) [8 µM], and port D hold rotenone [10 µM] and antimycin A [5 µM]. Six measurement cycles were performed prior to the first injection. Following the injection oligomycin A, four measurement cycles were conducted. Whereas subsequent injections were reduced to three measurement cycles.

### Mouse interleukin-6 (IL-6) enzyme-linked immunosorbent assay (ELISA)

The enzyme-linked immunosorbent assay (ELISA) was performed following the manufacturer’s instruction of BioLegend’s ELISA MAX™ Deluxe Set (BioLegend). The medium samples were diluted 1:5 with 1 × assay diluent A and analyzed as duplicates. As a stop solution, 2 N H_2_SO_4_ was used, and afterward, the absorbance at 450 nm and 570 nm was measured with the Tecan Infinite^®^ 200 Spark^®^. The protein concentration was determined through the linear standard curve of the measured standards and data was normalized to cell protein amount or cell number.

### Bicinchoninic acid protein assay

For the BCA protein assay, the Pierce™ BCA Protein Assay Kit (Cat.# 23250, Thermo Scientific) was used and performed according to the manufacturer’s instructions to normalize the samples to the protein concentration. Samples used for GC-MS measurement were dried in a 96-well cell culture plate before dissolving in 1× radioimmunoprecipitation assay (RIPA) buffer (Cat.# 20-188, Merck). The BCA protein assay for the Seahorse assay was performed in the cell culture plate. The medium was removed, and the cells were lysed in 1× RIPA buffer.

### Quantitative real-time polymerase chain reaction (qRT-PCR)

The RNA isolation was performed according to the manufacturer’s instructions for the NucleoSpin^®^ RNA Kit (Cat.# 740955.250, Machery-Nagel). RNA was isolated from a 12-well cell culture plate. Before lysing the cells, the supernatant was collected in 1.5 m1 reaction tubes and centrifuged at 300 × *g* at 4 °C for 5 min. The medium was stored at -20 °C for further use. The RNA was eluted with 40 µl of RNase-free H_2_O. The concentration measurement was performed with the Tecan Infinite^®^ 200 Spark^®^. 10 µl RNA (50 ng/µl) per sample was reverse transcribed into cDNA using High-Capacity cDNA Reverse Transcriptase kit (Cat.# 4368813, Thermo Fisher Scientific). Individual 10 μl SYBR Green real-time PCR reactions consisted of 2 μl of diluted cDNA, 5 μl iTaq Universal SYBR Green Supermix (Cat.# 1725124, BIO-RAD), and 1.5 μl of each 10 μM forward and reverse primers. For standardization of quantification, *60S ribosomal protein* (*Rpl27*) was amplified simultaneously. PCR was carried out in 96-well plates (MicroAmp™ Optical 96-Well Reaction Plate, Cat.# N8010560, Applied Biosystems by Life technologies®) and centrifuged at 300 x g for 30 sec before being measured with QuantStudio Design & Analysis Software v1.5.2 (Thermo Fisher Scientific).

The following primers were used. *Irg1* forward: GCAACATGATGCTCAAGTCTG; *Irg1* reverse: TGCTCCTCCGAATGATACCA; *Rpl27* forward: GCGATCCAAGATCAAGTCCTTTG; *Rpl27* reverse: TCAAAGCTGGGTCCCTGAACAC; *Slc13a3* forward: CAGGGAAATGGACTGCGAACAG; *Slc13a3* reverse: TCCTCCTTGGAGTCATCAGGCA;

### Statistics

Data visualization and statistical analysis were performed using GraphPad Prism (v10.3.1, GraphPad Software), Adobe Illustrator CS6 (v24.1.2, Adobe Inc.), and Biorender.com. The type and number of replicates and the statistical test used are described in each figure legend. Data are presented as means ± standard error of mean (SEM), box (25^th^–75^th^ percentile with median line) and whiskers (minimum to maximum values), or truncated violin plot. Experiments were independently repeated as indicated in the figure legend. *P* values were calculated using an unpaired *t*-test, one-way *ANOVA* or two-way *ANOVA* with Fisher’s least significant difference (LSD) post hoc test. For all tests, *p* < 0.05 was considered significant with \**p* < 0.05; \*\**p* < 0.01; \*\*\**p* < 0.001, and \*\*\*\**p* < 0.0001 as indicated in each figure legend.

## Results

### Alternative carbon sources influence immune metabolism in macrophages

To investigate how alternative carbon sources impact macrophage metabolism and function, we cultured RAW264.7 macrophage-like cells in standard DMEM (Ctr) or supplemented with alanine (ala), β-hydroxybutyrate (βHB), or lactate (lac) at concentrations approximating physiological plasma levels. These substrates are absent from standard DMEM but are present in plasma and serve as alternative carbon sources fueling the acetyl-Coenzyme A pool for TCA cycle metabolism and citrate synthesis^3,7^. Exposure to alanine, βHB, or lactate did not significantly influence LPS-induced IL-6 secretion indicating that these alternative carbon sources may not perturb inflammatory activation (Fig. 1a). Likewise, *Irg1* expression was upregulated upon pro-inflammatory stimulation but remained unaffected by the presence of alternative carbon sources (Fig. 1b). LPS treatments increased itaconate levels across all conditions, with slightly elevated itaconate levels in βHB- and lactate-treated cells compared to control conditions (Fig. 1c). We also quantified expression of the plasma membrane transporter *Slc13a3* which facilitates the transport of citrate, succinate, and itaconate^26–28,30^. We observed a significant downregulation of *Slc13a3* expression in response to LPS stimulation independent of exogenous carbon source supplementation (Fig. 1d). These data suggest that while macrophages retain their inflammatory response in cultures with alternative carbon source supplementation, expression of *Slc13a3* is downregulated during immune responses in our model system. Our standard high-glucose DMEM contains 25 mM glucose and 4 mM glutamine which are the primary carbon sources for central carbon metabolism. Thus, we applied stable isotope tracing and mass spectrometry approaches and cultured RAW264.7 cells in the presence of [U-^13^C_6_]glucose or [U-^13^C_5_]glutamine and addition of alanine, βHB or lactate (Fig. 1e). We observed increased cellular abundances of these alternative carbon sources indicating that cells take them up from the medium (Fig. S1a, b). Supplementation with alanine and lactate, which are converted into pyruvate, resulted in reduced incorporation of ^13^C from glucose into pyruvate, alanine, and lactate indicating dilution effects. These patterns were observed in both LPS-treated and untreated cultures suggesting a shared metabolic response to exogenous carbon sources (Fig. 1f, Fig. S1c). Similarly, labeling on TCA cycle intermediates was slightly decreased suggesting the use of these alternative carbon sources for TCA cycle metabolism (Fig. 1f). Since this utilization may impact other carbon sources, we quantified incorporation of [U-^13^C_5_] glutamine-derived carbons into TCA cycle metabolism. We observed that lactate, βHB, and alanine reduced ^13^C labeling on TCA cycle intermediates with alanine supplementation having the highest impact (Fig. 1g, Fig. S1d). Our tracing experiments further revealed reduced M1 labeling on itaconate from ^13^C glucose, and decreased M4 labeling from ^13^C glutamine in cells cultured in the presence of alanine, βHB, or lactate (Fig. 1h, I, Fig. S1e, f). These labeling patterns support the biosynthesis pathway of itaconate through decarboxylation of cis-aconitate catalyzed by ACOD1 (Fig. S1g, h, Fig. 1e). These data suggest a diminished reliance on glucose and glutamine as a carbon source when exogenous alternative carbon sources are present. Our data further support the metabolic flexibility of macrophages to rewire central carbon metabolism for itaconate synthesis (Fig. 1j). Thus, in our culture models, cells are metabolically adaptable and incorporate physiologically relevant alternative carbon sources without compromising LPS-induced inflammatory responses.

**Figure 1:**
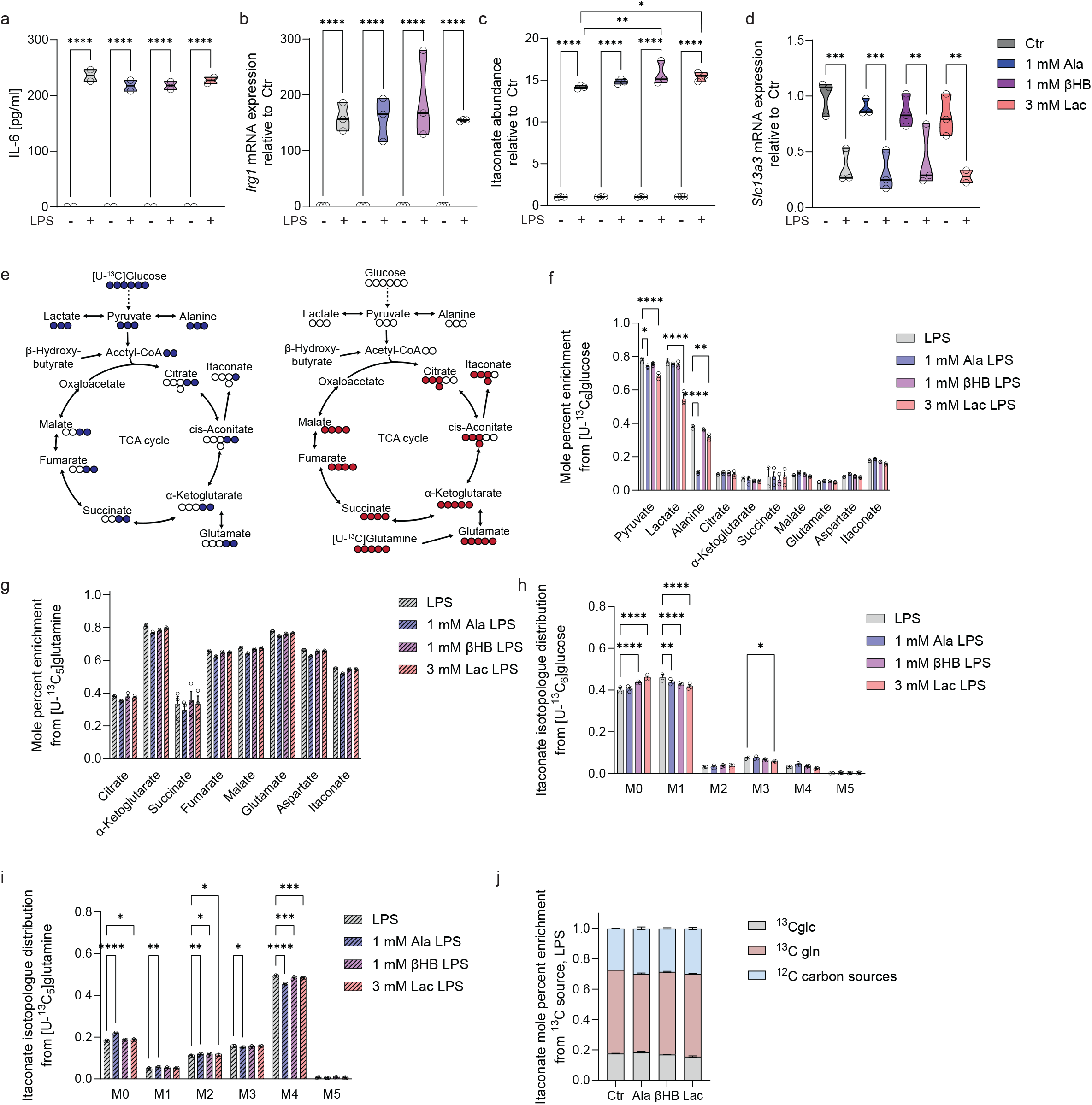
Effects of alternative carbon sources on TCA cycle metabolism and immune responses in macrophages. **a**, IL-6 concentration in different media. **b**, Expression level of *Irg1* in different media. **c**, Itaconate abundances in different media. **d**, Expression level of *Slc13a3* in different media. **e**, Metabolic map depicting atom transitions from [U-^13^C_5_]glutamine (red) and [U-^13^C_6_]glucose (blue). Open circles depict ^12^C, closed circles ^13^C atoms. **f**, Mole percent enrichment on metabolites from [U-^13^C_6_]glucose in different media. **g**, Mole percent enrichment on metabolites from [U-^13^C_5_]glutamine in different media. **h**, Isotopologue distribution on itaconate from [U-^13^C_6_]glucose. **i**, Isotopologue distribution on itaconate from [U-^13^C_5_]glutamine. **j**, Mole percent enrichment on citrate and itaconate from different tracers. RAW264.7 cells were treated with 10 ng/ml LPS for 6 h in DMEM supplemented with 1 mM alanine (ala), 1 mM β-hydroxybutyrate (βHB), or 3 mM lactate (lac) and results are compared to control conditions. Data are presented as truncated violin blot, (a-d), bar plots ± SEM (f-i) or stacked bar plots ± SEM (j) obtained from three cellular replicates. *P*-values were calculated by one-way *ANOVA* (a-d) or two-way *ANOVA* (f-i) with \**p* < 0.05, \*\**p* < 0.01, \*\*\**p* < 0.001, \*\*\*\**p* < 0.0001

### Modified culture media with alternative carbon source availability alter immune responses in macrophages

Since we observed minor effects on immune responses in cultures supplemented with alternative carbon sources in standard DMEM, we modified the standard culture medium to better reflect physiological conditions by creating a custom medium, referred to as modified DMEM (mDMEM) (Fig. 2a, b)^7^. To better mimic physiological nutrient conditions and investigate potential immunoregulatory effects, we reduced glucose concentrations from 25 mM to 5 mM, and glutamine concentrations from 4 mM to 0.5 mM closely approximating plasma levels. In addition to lowering these primary carbon sources, we supplemented mDMEM with alternative carbon substrates including acetate, alanine, asparagine, citrate, lactate, pyruvate, and βHB. These substrates are absent from standard DMEM but are detectable in plasma and known to be oxidized in mice (Fig 2a, b)^6,7^. This modification resulted in a 47.4 % reduction in total carbon source availability compared to standard DMEM (Fig. 2c). To maintain nutrient availability and mitigate substrate depletion, specifically of glutamine, we performed regular medium changes during our culture experiments (Fig. 2d).

**Figure 2:**
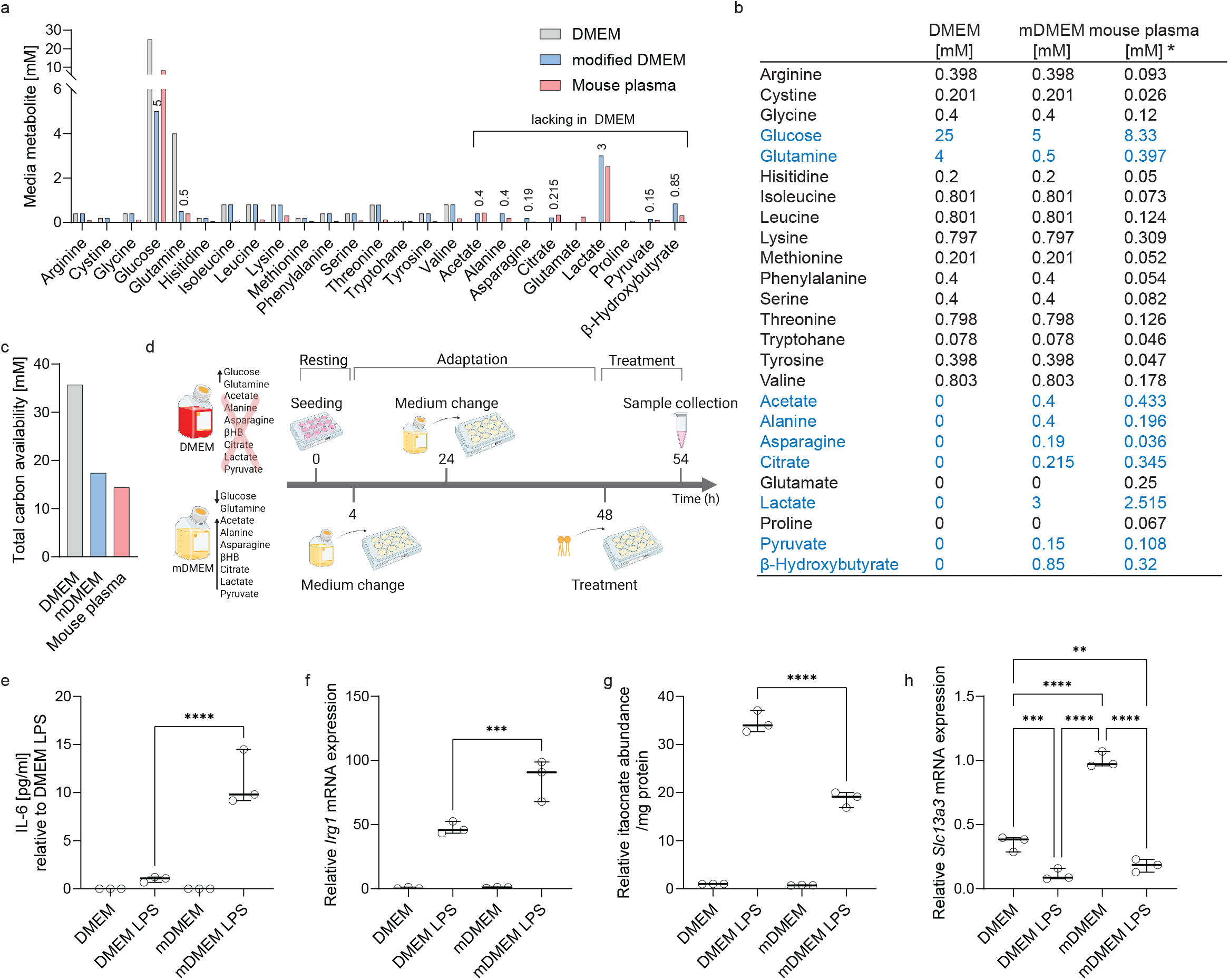
Modified culture media with alternative carbon source availability alter immune responses in macrophages. **a**, Media composition of DMEM (grey), compared to modified DMEM (mDMEM, blue) and mouse plasma (red), starting with the similar concentrations on the left side of the x-axis and the differing components on the right. **b**, Formulation of DMEM and mDMEM compared to mouse plasma (mouse plasma data taken from Sugimoto *et al*., 2012, Hui *et al*., 2017, and Kaymak *et al*., 2022). **c**, Total carbon availability in media. **d**, Schematic depicting the experimental overview. Cells were incubated for 4 h after seeding before changing to mDMEM. Media was changed after 36 h of culture. Tracer media and LPS were added at 48 h for 6 h before sample collection, created with Biorender.com. **e**, IL-6 concentration in media. **f**, Expression level of *Irg1*. **g**, Itaconate abundances. **h**, Expression level of *Slc13a3*. RAW264.7 cells were adapted for 44 h to the modified DMEM (mDMEM) before 6 h treatment with 10 ng/ml LPS. Data are presented as bar plots (a, c) or box (25^th^–75^th^ percentile with median line) and whiskers (minimum to maximum values) obtained from three cellular replicates. Each experiment was repeated independently two (e, f, h) or three (g) times. *P*-values were calculated by one-way *ANOVA* (e-h) with \**p* < 0.05, \*\**p* < 0.01, \*\*\**p* < 0.001, \*\*\*\**p* < 0.0001

We cultured RAW264.7 cells in either standard DMEM or mDMEM and observed increased IL-6 secretion in cells cultured in mDMEM (Fig. 2e) indicating enhanced inflammatory responses under more physiological nutrient conditions. Further, expression of *Irg1* was increased in mDMEM while itaconate levels were reduced relative to cells in standard DMEM (Fig. 2f, g).

Next, we analyzed the expression of *Slc13a3* and observed decreased *Slc13a3* expression levels in response to LPS in both media (Fig. 2h). Levels of *Slc13a3* was significantly higher in mDMEM cultures compared to standard DMEM cultures under unstimulated conditions suggesting that the nutrient microenvironment may influence transporter expression. Collectively, these results demonstrate that media with reduced glucose and glutamine concentrations and additional alternative carbon sources (mDMEM) influence macrophage biology in our model system.

### Physiological carbon sources induce metabolic reprogramming in mDMEM

To assess whether the reduced total carbon content in mDMEM alters macrophage metabolism, we traced central carbon metabolism using [U-^13^C_6_]glucose and [U-^13^C_5_]glutamine. We observed decreased intracellular abundances of TCA cycle intermediates, specifically αKG and malate, in mDMEM compared to standard DMEM cultures (Fig. 3a). Intracellular alanine and aspartate levels were elevated reflecting their supplementation in the mDMEM formulation (Fig. 3b). These changes occurred independently of the pro-inflammatory activation state of the macrophages. Labeling from [U-^13^C_6_]glucose on pyruvate, citrate, and alanine in cells cultivated in mDMEM was reduced consistent with dilution from the corresponding unlabeled supplemented metabolites (Fig. 3c, Fig. S2a-f). Next, we applied [U-^13^C_5_]glutamine and observed reduced labeling on TCA cycle intermediates in mDMEM cultures (Fig. 3d). Labeling of itaconate and citrate was also reduced in mDMEM following both tracers (Fig. 3e, f, Fig. S2g).

**Figure 3:**
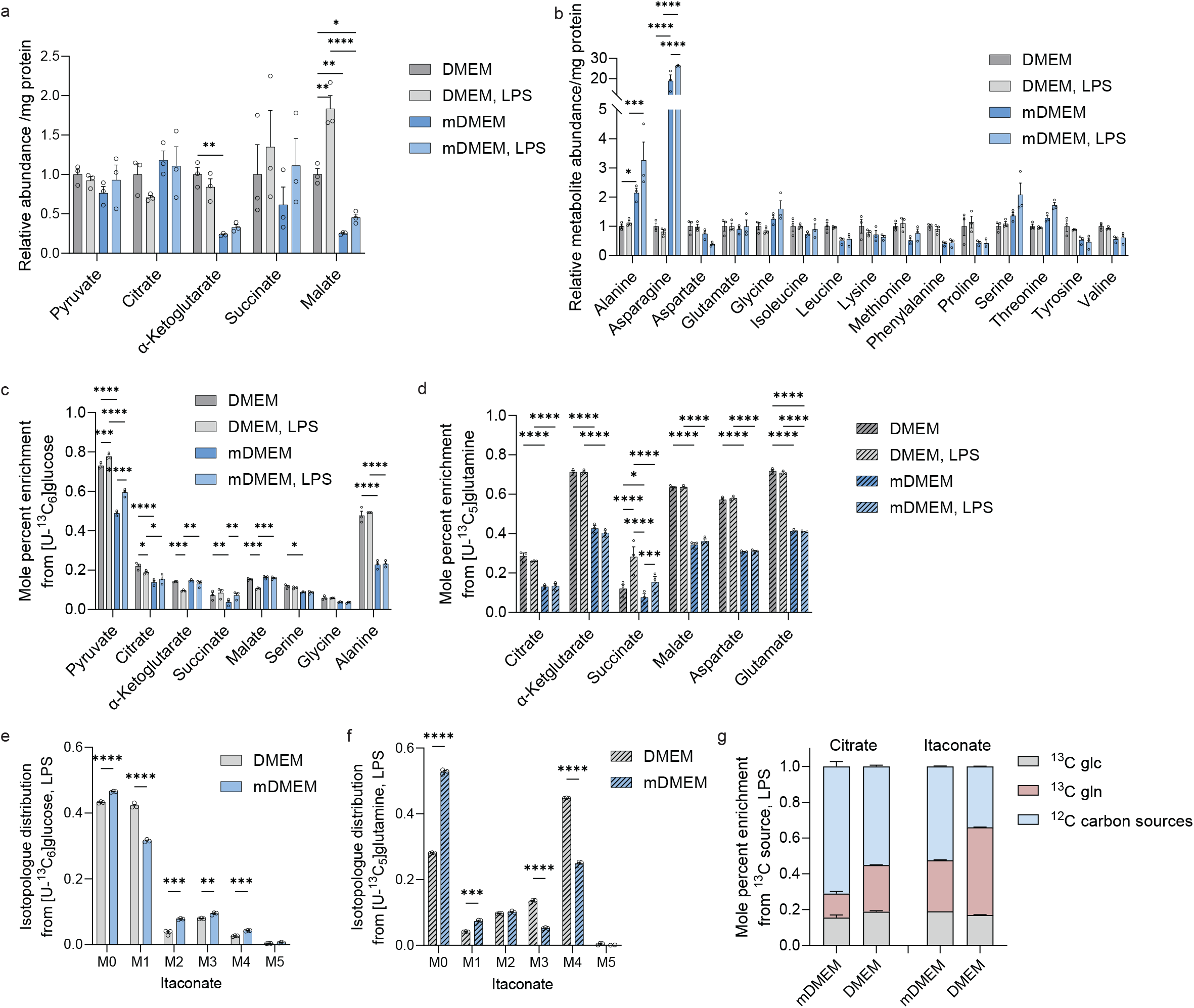
Physiological carbon sources induce metabolic reprogramming. **a**, Metabolite abundances in cultures with DMEM compared to mDMEM. **b**, Amino acid abundances in cultures with DMEM compared to mDMEM. **c**, Mole percent enrichment on metabolites from [U-^13^C_6_]glucose in different media. **d**, Mole percent enrichment on metabolites from [U-^13^C_5_]glutamine in different media. e, Isotopologue distribution on itaconate from [U-^13^C_6_]glucose. **f**, Isotopologue distribution on itaconate from [U-^13^C_5_]glutamine. **g**, Mole percent enrichment on citrate and itaconate from different labeled carbon sources. RAW264.7 cells were adapted to the modified DMEM (mDMEM) 44 h before a 6 h treatment with 10 ng/ml LPS. Data are presented as bar plots or stacked bars ± SEM obtained from three cellular replicates. Each experiment was repeated independently two (c-g) or three (a, b) times. *P*-values were calculated by two-way *ANOVA* (a-d) or multiple unpaired t-test (e, f) with \**p* < 0.05, \*\**p* < 0.01, \*\*\**p* < 0.001, \*\*\*\**p* < 0.0001.

Together, these data indicate that altered nutrient composition of mDMEM decreases the contribution of glucose and glutamine to TCA cycle metabolism (Fig. 3g). The reduced tracer incorporation suggests that one or more supplemented metabolites in mDMEM may alter the use of glucose and glutamine thereby affecting substrate routing into mitochondrial metabolic pathways.

### Citrate influences immune responses in media with alternative carbon sources

Our labeling experiment revealed decreased utilization of glucose and glutamine for TCA cycle metabolism and specifically on citrate in mDMEM cultures (Fig. 3). Since we also observed changes in *Slc13a3* expression we tested whether extracellular citrate contributed to these effects. Thus, we cultured RAW264.7 cells in mDMEM with and without supplementing 0.215 mM unlabeled citrate in the presence of [U-^13^C_5_]glutamine (Fig. 4a). This concentration closely matches the concentrations found in plasma and present in mDMEM. Citrate supplementation reduced labeling on citrate consistent with isotopic dilution from the exogenous citrate pool (Fig. 4b, c). However, labeling on itaconate and other TCA cycle intermediates was unaffected suggesting that citrate did not broadly affect glutamine flux into downstream TCA cycle intermediates (Fig. 4b-d). To directly test whether cells metabolize exogenous citrate, we supplemented 0.215 mM [2,4-^13^C_2_]citrate tracer to mDMEM cultures (Fig. S3a, S3b)^31^. Itaconate and other TCA cycle intermediates were unlabeled indicating minimal oxidation of exogenous citrate via the TCA cycle under these conditions (Fig. S3c).

**Figure 4:**
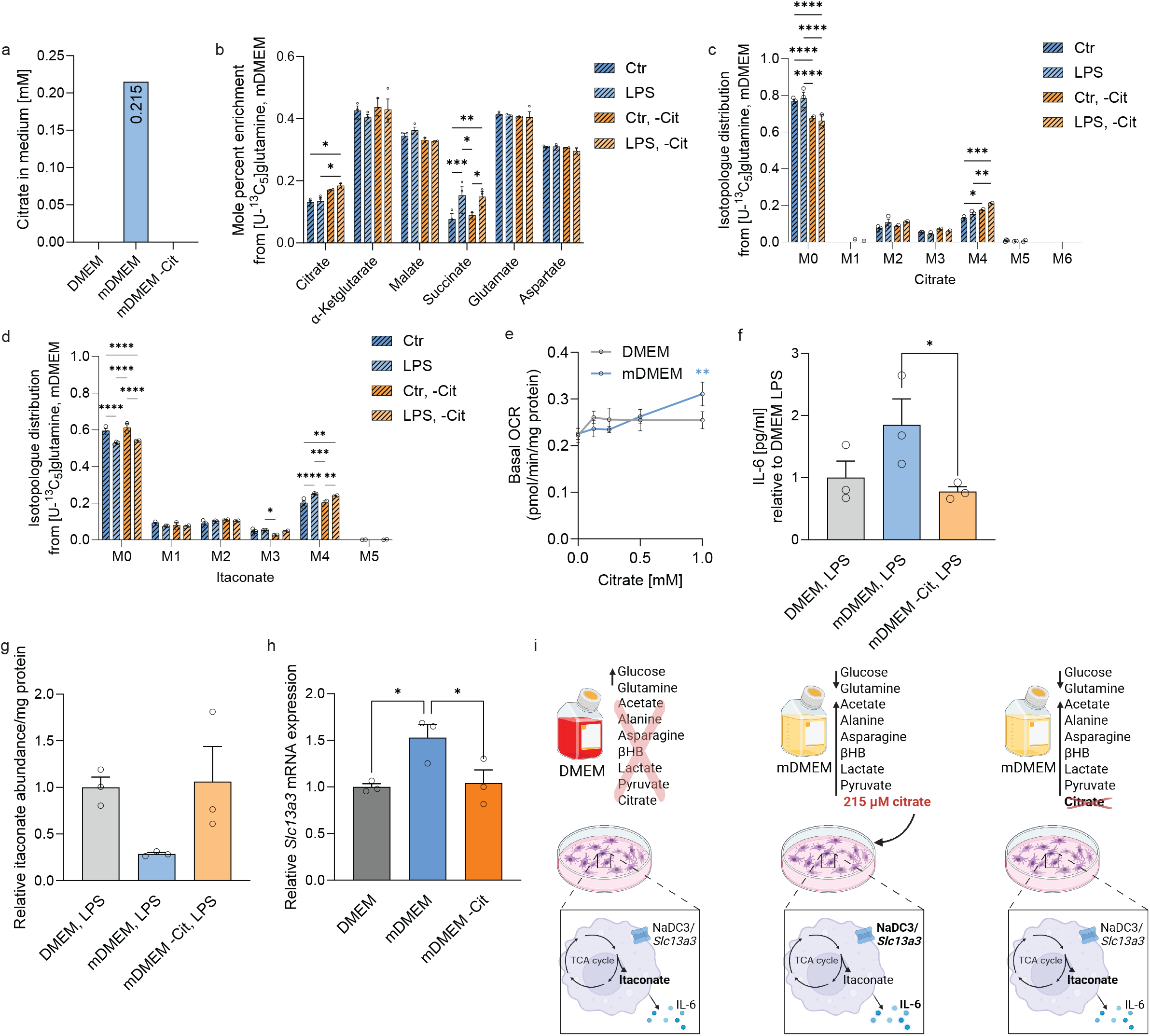
Citrate influences immune responses in media with alternative carbon sources. **a**, Citrate concentration in different culture media. **b**, Mole percent enrichment from [U-^13^C_5_]glutamine on metabolites in cells cultivated in mDMEM with and without citrate. **c**, Isotopologue distribution of citrate from [U-^13^C_5_]glutamine. **d**, Isotopologue distribution of itaconate from [U-^13^C_5_]glutamine. **e**, Basal oxygen consumption rate in cells cultured in DMEM and adapted DMEM with increasing citrate concentrations. **f**, IL-6 concentration in media in response to LPS. **g**, Itaconate abundances in response to LPS. **h**, Expression level of *Slc13a3*. **i**, Schematic overview illustrating effects of media composition on immunometabolism, created with Biorender.com. RAW264.7 cells were adapted to the modified DMEM (mDMEM) with and without citrate supplementation 44 h before a 6 h treatment with 10 ng/ml LPS. Data are presented as bar plots ± SEM obtained from three cellular replicates (b-d, f-h) or points with connecting lines ± SEM obtained from six cellular replicates (e). Each experiment was repeated independently two times. *P*-values were calculated by two-way *ANOVA* (b-e) or one-way *ANOVA* (f-h) with \**p* < 0.05, \*\**p* < 0.01, \*\*\**p* < 0.001, \*\*\*\**p* < 0.0001. Significance in e is depicted compared to 0 mM citrate.

Next, we quantified the effects of increasing citrate levels on mitochondrial respiration and observed that citrate did not alter oxygen consumption rates (OCR) in standard DMEM medium (Fig. 4e). In contrast, higher citrate concentrations in mDMEM increased OCR. These findings suggest that the influence of citrate on mitochondrial respiration is dependent on the cellular environment.

Given the observed changes in mitochondrial respiration, we analyzed functional changes in macrophages cultured in the modified DMEM (mDMEM) with or without citrate supplementation. We observed that LPS-induced secretion of IL-6 was significantly increased in mDMEM compared to standard DMEM cultures (Fig. 4f). Of note, removal of citrate from mDMEM cultures restored IL-6 secretion to levels comparable to those observed in standard DMEM cultures (Fig. 4f). Further, itaconate accumulation was reduced in mDMEM cultures and partly restored upon citrate removal (Fig. 4g). We also observed that removal of citrate from mDMEM cultures reduced the expression of *Slc13a3* almost to that of standard DMEM cultures (Fig. 4h). These data indicate that physiologically relevant carbon sources modulate immune responses and immunometabolism in our culture models and that citrate plays a role in these processes (Fig. 4i).

## Discussion

Our study demonstrates that culturing macrophages in a physiologically relevant medium (mDMEM) with reduced glucose and glutamine concentrations and supplemented with alternative carbon sources, including citrate, alters cellular metabolism and immune responses. Our metabolic flux analysis suggests a shift toward these supplemented substrates and identifies that citrate acts as a modulator of macrophage immunometabolism.

Our metabolic flux analyses suggest that macrophages exhibit metabolic flexibility and reroute carbon flux through the TCA cycle by using alternative substrates when canonical sources such as glucose and glutamine are limited. These substrates fuel mitochondrial metabolism and synthesis of immunomodulatory metabolites such as itaconate which accumulates to millimolar levels during immune responses. We identify that citrate is not a direct carbon fuel for TCA cycle metabolism but modulates immunometabolic pathways. Citrate supplementation at physiological levels of 0.215 mM increased pro-inflammatory IL-6 secretion while reducing intracellular itaconate^32,33^. Thus, citrate may shape the balance between inflammatory signaling and metabolic intermediates for immunoregulation^13,34,35^. The presence of physiological carbon sources also modulated effector function of CD8+T cells by shifting carbon utilization away from glucose toward lactate as a biosynthetic fuel^7^. This suggests that systemic nutrient availability may regulate macrophage metabolism, thereby modulating immune responses and itaconate metabolism. Further, under more physiologically relevant conditions, citrate acts as a metabolic signal that shapes the balance between inflammatory responses and central carbon metabolism.

Citrate supplementation induced expression of plasma membrane transporter *Slc13a3* (NaDC3). This transcriptional adaptation may influence nutrient uptake in response to environmental changes and may facilitate metabolic flexibility. Since *Slc13a3* is also involved in itaconate transport^26–28^, our findings may be relevant to understanding the dynamic regulation of itaconate metabolism^36^. Exclusion of citrate from the medium reversed these effects highlighting its role as a metabolic signal that modulates immunological phenotype of macrophages^13,37^. While citrate is abundant in plasma and tissues, and high levels are linked to SLC13A5 citrate transport disorder^32,33,38^, it is absent from most standard culture media^39^. Consequently, potential interactions between exogenous citrate and itaconate metabolism may have been overlooked due to the lack of physiologically relevant citrate levels in in vitro culture models.

This study focuses on alternative carbon sources that are absent from standard DMEM, namely acetate, alanine, asparagine, citrate, lactate, pyruvate, and βHB^7^. Future investigations may consider examining the impact of varying concentrations of other DMEM metabolites, such as amino acids, as these were not adjusted to physiologically relevant levels in our experiments. Further studies may also elucidate the impact of the individual physiologically relevant carbon sources as well as the molecular mechanisms by which citrate modulates immune responses and the subsequent effects on metabolic networks^30^.

Collectively, our findings highlight that media composition and tissue microenvironments are dynamic variables that influence macrophage biology and immunometabolism where citrate plays a role. The ability to adapt carbon source usage enables macrophages to preserve TCA function, itaconate production, and immune responses across changing nutrient landscapes. Our study emphasizes the need to understand metabolic rewiring in response to physiologically relevant nutrients and tailoring metabolic interventions during immune responses.

## Acknowledgment

We thank all members of the Cordes Lab for helpful discussions. This study was supported, in part, by a Cambridge Isotope Laboratories Inc. (CIL) Grant as well as by internal funds from the Helmholtz Centre for Infection Research (to T.C.) and Technische Universität Braunschweig (to T.C.). We acknowledge support from the Open Access Publication Funds of Technische Universität Braunschweig.

## Availability of data

All of the data associated with the study are in the manuscript. Source data will be deposited in the repository platform of Technische Universität Braunschweig with a dedicated doi upon acceptance of this manuscript.

## Contributions

T.C. and L.W. conceived and designed the study. L.W. and H.F.W. performed experiments. T.C., L.W., and H.F.W. analyzed the data. T.C. and L.W. wrote the manuscript with input from all authors.

## NList of abbreviations

^12^C: Carbon-12 (unlabeled)
^13^C: arbon-13 (labeled)
Ala: Alanine
BCA: Bicinchoninic acid
DMEM: Dulbecco’s modified Eagle’s medium
mDMEM: Modified Dulbecco’s modified Eagle’s medium
ELISA: Enzyme-linked immunosorbent assay
FBS: Fetal bovine serum
FCCP: Trifluoromethoxy carbonylcyanide phenylhydrazone
GC-MS: Gas chromatography-mass spectrometry
IL-6: Interleukin-6
IRG1/ACOD1: Immune responsive gene 1 protein / cis-aconitate dehydrogenase 1
*Irg1;*: Immunoresponsive gene 1
Lac: Lactate
LPS: Lipopolysaccharide
LSD: Fisher’s least significant difference
MPE: Mole percent enrichment
NAD: Nicotinamide adenine dinucleotide
NaDC3/SLC13A3: Sodium-dependent dicarboxylate transporter 3 protein
OCR: Oxygen consumption rate
P/S: Penicillin/streptomycin
qRT-PCR: Quantitative real-time polymerase chain reaction
RIPA: Radioimmunoprecipitation assay
*Rpl27;*: 60S ribosomal protein
SDH: Succinate dehydrogenase
SEM: Standard error of mean
*Slc13a3;*: Solute carrier family 13 member 3 gene
TCA: Tricarboxylic acid
αKG: α-Ketoglutarate
βHB: β-Hydroxybutyrate

---

**Supplementary Figure 1:**
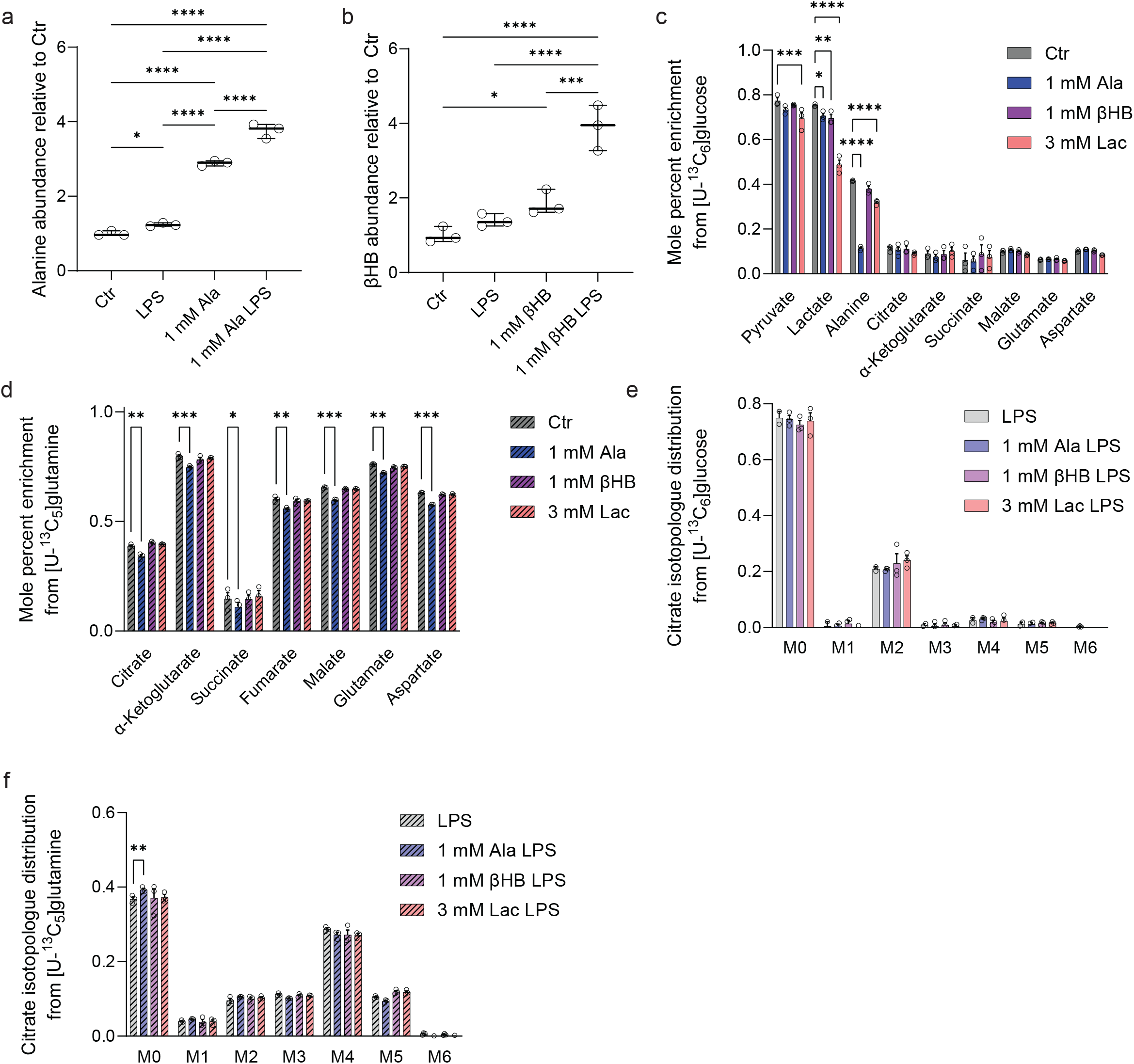
Physiological carbon sources influence mitochondrial metabolism. **a**, Alanine abundance in cultures with exogenous alanine. **b**, ß-Hydroxybutyrate (βHB) abundance in cultures with exogenous βHB. **c**, Mole percent enrichment on metabolites from [U-^13^C_6_]glucose in different media. **d**, Mole percent enrichment on metabolites from [U-^13^C_5_]glutamine in different media. **e**, Isotopologue distribution on citrate from [U-^13^C_6_]glucose. **f**, Isotopologue distribution on citrate from [U-^13^C_5_]glutamine. RAW264.7 cells were treated with 10 ng/ml LPS for 6 h in DMEM supplemented with 1 mM alanine (ala), 1 mM ß-hydroxybutyrate (βHB), or 3 mM lactate (lac) where indicated. Data are presented as box (25^th^– 75^th^ percentile with median line) and whiskers (minimum to maximum values) (a, b) or bar plots ± SEM (c-h) obtained from three cellular replicates. Each experiment was repeated independently two times (a, b). *P*-values were calculated by one-way *ANOVA* (a, b), unpaired *t*-test (c, d) or two-way *ANOVA* (e-h) with \**p* < 0.05, \*\**p* < 0.01, \*\*\**p* < 0.001, \*\*\*\**p* < 0.0001

**Supplementary Figure 2:**
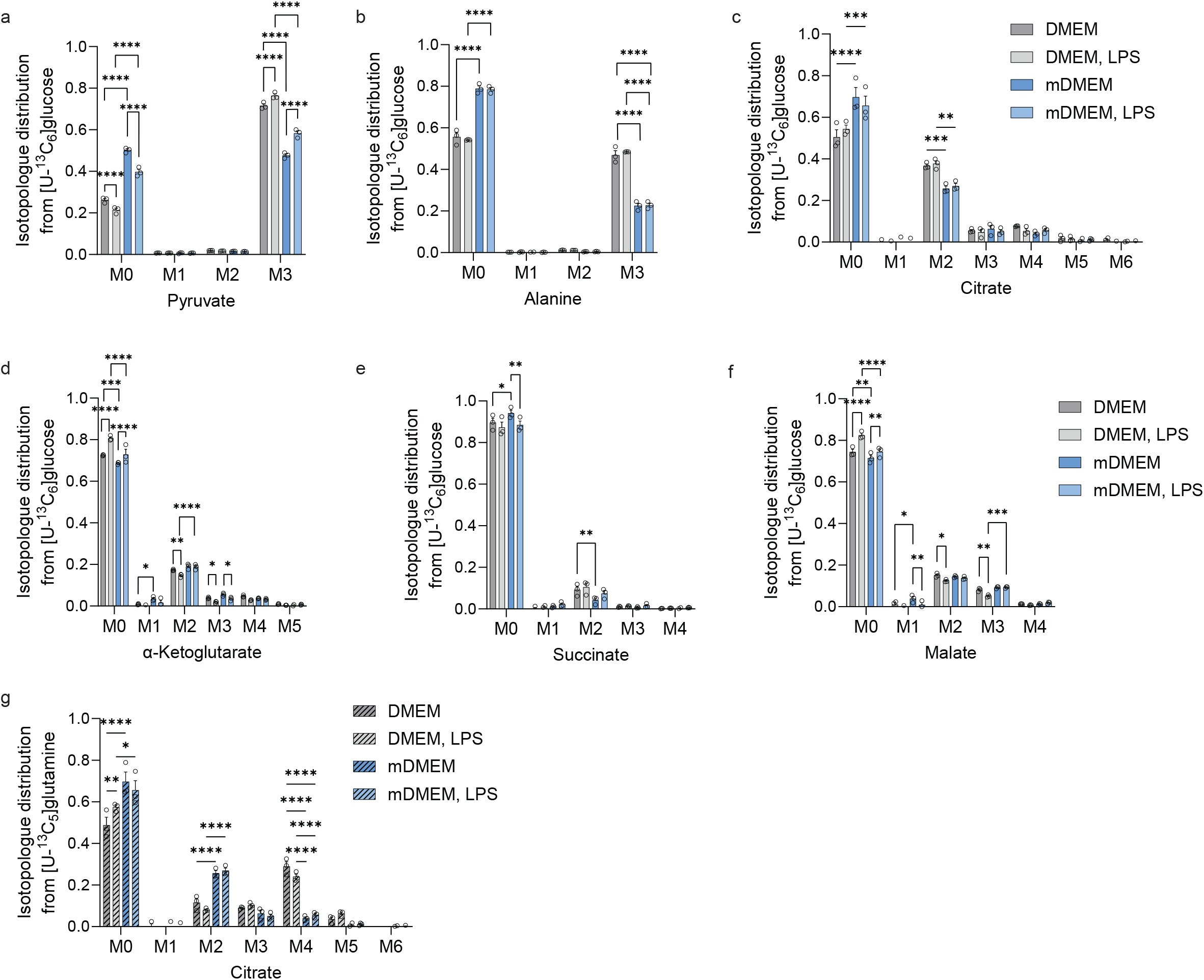
Alternative carbon sources influence glucose and glutamine utilization. Isotopologue distribution from [U-^13^C_6_]glucose on **a**, pyruvate, **b**, alanine, **c**, citrate, **d**, αKG, **e**, succinate, and **f**, malate. Isotopologue distribution from [U-^13^C_5_]glutamine on citrate. RAW264.7 cells were adapted to the modified DMEM (mDMEM) 44 h before a 6 h treatment with 10 ng/ml LPS. Data are presented as bar plots ± SEM obtained from three cellular replicates. Each experiment was repeated independently two times. P-values were calculated by two-way *ANOVA* with *p < 0.05, **p < 0.01, ***p < 0.001, ****p < 0.0001.

**Supplementary Figure 3:**
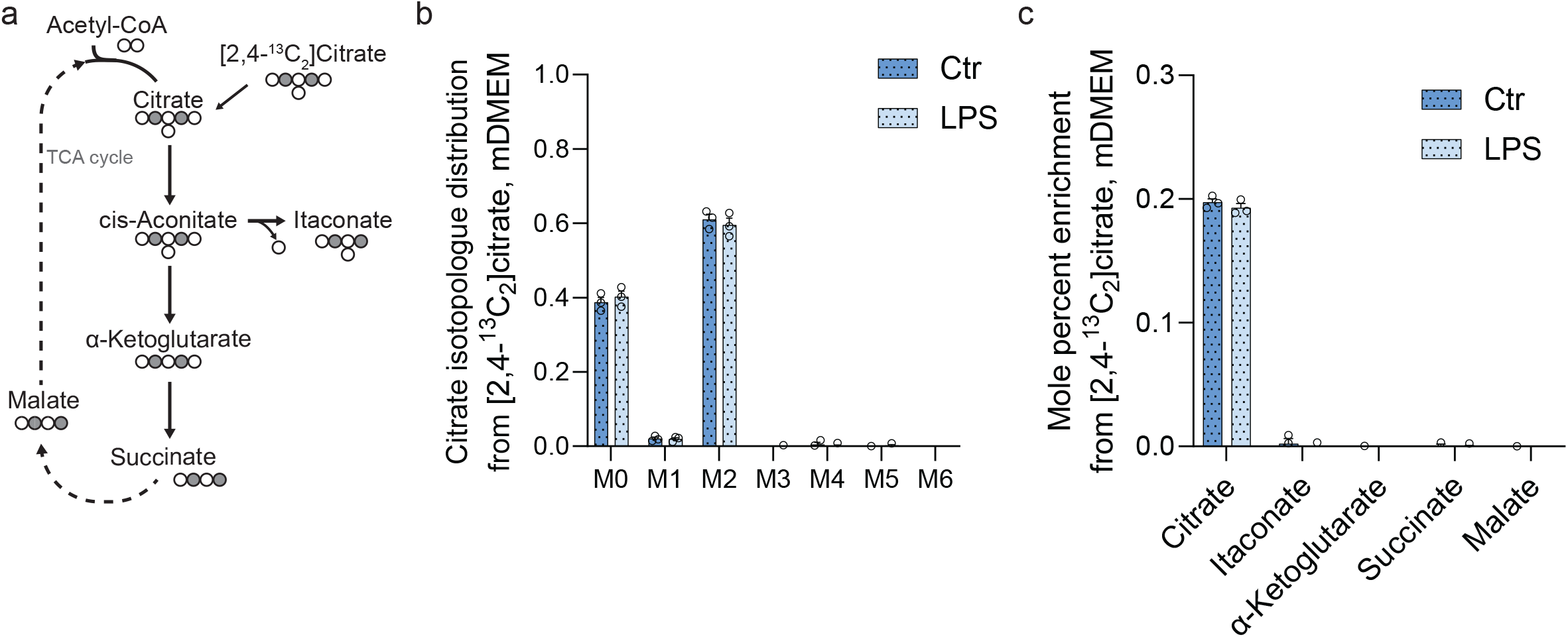
Citrate trace into TCA cycle metabolism. **a**, Metabolic map depicting atom transitions from [2,4-^13^C_2_]citrate. Open circles depict ^12^C, closed circles ^13^C atoms. **b**, Isotopologue distribution from 0.215 mM [2,4-^13^C_2_]citrate on citrate. **c**, Mole percent enrichment on TCA cycle intermediates from [2,4-^13^C_2_]citrate. RAW264.7 cells were adapted to the modified DMEM (mDMEM) with and without citrate supplementation 44 h before a 6 h treatment with 10 ng/ml LPS. Data are presented as bar plots ± SEM obtained from three cellular replicates.

